# Improving the application of Important Plant Areas to conserve threatened habitats: a case study of Uganda

**DOI:** 10.1101/2024.01.04.572590

**Authors:** Sophie L. Richards, James Kalema, Samuel Ojelel, Jenny Williams, Iain Darbyshire

**Author notes:** Corresponding author: Sophie L. Richards.

## Abstract

Important Plant Areas (IPAs) are a successful method of identifying priority areas for plant conservation. Assessment of IPAs, however, often relies on criteria related to species, while incorporation of habitats has been less consistent. Using Uganda as a case study, we test the application of the threatened habitat criterion – criterion C. We identified nationally threatened habitats using Red List of Ecosystems criteria and assess, for the first time, how differing application of thresholds under criterion C can influence IPA network outcomes. Eleven threatened habitats were identified, with declines switching from predominantly forest to savanna after the mid-20^th^ century. Significantly, we found current IPA guidance on use of Criterion C needlessly limits the number of sites that qualify as IPAs. The “five best sites” threshold is reserved for countries where quantitative data is unavailable, however, the application of the relevant thresholds to quantitative data largely generated fewer than five IPAs, comparably limiting conservation opportunities identified. We recommend, therefore, that the “five best” threshold is available for application on both qualitative and quantitative data. This will bolster the value of IPAs in conserving and restoring threatened and ecologically important habitats under the Kunming-Montreal Global Biodiversity Framework.

## Introduction

Ecosystems, encompassing species communities, the genetic diversity and physical environment they exist within, are complex components of biodiversity which contribute immeasurably to human wellbeing through the provision of ecosystem services (IPBES, 2019; Keith et al., 2023). Protecting ecosystems from anthropogenic perturbations will also help conserve rare and threatened species that exist within (Bland et al., 2017). However, 75% of Earth’s land surface is significantly altered with many ecosystems becoming more homogenised, losing both biodiversity and ecosystem services (IPBES, 2019).

The Kunming-Montreal Global Biodiversity Framework (CBD, 2022) commits signatories to maintain, enhance, or restore the integrity, connectivity and resilience of all ecosystems and substantially increase the area of natural ecosystems by 2050 (Goal A). Parties must also ensure that by 2030, 30% of degraded ecosystems are under effective restoration (Target 2) and an ecologically representative 30% of areas important for biodiversity and ecosystem function are effectively conserved and managed through protected areas (Target 3).

Identification of the most threatened ecosystems is a key step towards prioritising conservation action and meeting these aims. Important Plant Areas (IPAs) are one way of undertaking such prioritisation of ecosystems.

IPAs are defined as the most important places in the world for wild plant and fungal diversity that can be protected and managed as specific sites (Anderson, 2002; Darbyshire et al., 2017). There are difficulties in defining ecosystems for conservation assessment as identified by previous authors (Boitani et al., 2015; Keith et al., 2013, 2015b), and therefore habitats or vegetation types are commonly used as a proxy for ecosystems within IPAs and other conservation prioritisations such as the IUCN Red List of Ecosystems (Bland et al., 2017; Keith et al., 2013).

Within the IPAs criteria, threatened habitats fall under criterion C. The process of applying this criterion has two stages: 1) identification of threatened habitats and 2) identification of sites that trigger IPAs based upon presence of these threatened habitats. Criterion C can be applied at different scales: globally under C(i), regionally under C(ii) and nationally under C(iii).

Despite this flexibility in application, criterion C has often been used secondarily to species-based criteria A and B. In Mozambique for example, all sites qualify under criterion A, based on threatened species, but just under half trigger criterion C and no site qualifies under C alone (Darbyshire et al., 2017).

Table 1 details how criterion C has been applied so far in the global tropics. The majority of these countries have included threatened habitats within assessments, largely under sub-criterion C(iii) (Box 1).

### Box 1

**IPA sub-criterion C(iii) (Darbyshire *et al*. 2017)**

Site contains nationally threatened or restricted habitat / vegetation type, AND/OR habitats that have severely declined in extent nationally.

Site known, thought or inferred to contain **≥10% of the national resource (area)** of the threatened habitat type OR site is among the best quality examples required to **collectively prioritise up to 20% of the national resource** OR the **5 “best sites”** nationally, whichever is the most appropriate.

Wherever possible, the national importance of the site should be documented by applying the threshold for the % of the national resource; the selection of ‘‘best sites’’ should only be applied where quantitative data are not available and cannot be inferred.

**Table 1:** Application of threatened habitats each tropical, or partially tropical, nation or territory with a completed IPA assessment according to the Plantlife database. ≥10%: site known, thought or inferred to contain ≥10% of the national resource (area) of the threatened habitat type. Cml. 20%: site is among the best quality examples required to collectively prioritise up to 20% of the national resource. 5 best: the five “best sites” nationally, whichever is the most appropriate. Asterisk refers to assessments carried out before the establishment of criterion C(iii) following Darbyshire et al. (2017), when assessment of threatened habitats was largely based on the European Habitats Directive and Bern list (Anderson, 2002).

| Country | Threatened | Method of threatened | Quantitative | Threshold used for IPA |  |  | Citation |
| --- | --- | --- | --- | --- | --- | --- | --- |
|  | habitats | habitat identification | data on | identification |  |  |  |
|  | assessed | (step one) | threatened | (step two) |  |  |  |
|  |  |  | habitat extent? | ≥10% | Cml. 20% | Five best |  |
| Bolivia<br>(Chiquitanía,<br>Santa Cruz) | Yes | Retrospective application of Red List of Ecosystems to IPAs | Generated post IPA-assessment | NA | NA | NA | (Martinez-Ugarteche et al., 2023) |
| British Virgin Islands | Yes, C(iii) | Extent of habitat and proof of continuous decline | Yes | No | No | Yes | (Barrios et al., 2019) |
| Cameroon | No | NA | NA | NA | NA | NA | (Murphy et al., 2023) |
| Ethiopia<br>(pilot) | Yes, C(iii) | Expert opinions | Yes | Yes | No | Yes | (House et al., 2023) |
| Guinea | Yes, C(iii) | Number of endemic and threatened species per habitat | Yes | Yes | No | No | (Couch et al., 2019) |
| Mozambique | Yes, C(iii) | Expert opinions | Yes | Yes | No | Yes | (Darbyshire et al., 2023) |
| Algeria<br>(North) | No* | NA | NA | NA | NA | NA | (Yahi et al., 2012) |
| Egypt | Yes* | Expert opinions | No | NA | NA | NA | (Shaltout & Eid, 2016) |

Criterion C is not prescriptive on how threatened habitats can be identified, to promote usability in countries with limited data. As a result, various strategies have been used to identify nationally threatened habitats (Table 1, stage one). Several rely on expert opinion rather than quantitative assessment of habitat. This may be due to a lack, or perceived lack, of data available to undertake these assessments.

Subsequent IPA selection (stage two) relies upon thresholds to capture sites which represent significant areas of the national resource (Box 1). IPA guidance states that the quantitative thresholds (site contains ≥10% of the national resource or site is among the best quality examples required to collectively prioritise up to 20% of the national resource) should always be applied unless only qualitative data on the extent of a habitat is available, in which case a site can be selected if it is believed to be one of the “five best” nationally (Darbyshire et al., 2017). However, the “five best” threshold has been applied to quantitative data in at least three IPA programmes (Table 1).

In this analysis, we evaluate both stages in the application of sub-criterion C(iii) using Uganda as a case study. At stage one, we apply Red List of Ecosystems (RLE) criteria to identify threatened habitats. The RLE was developed to identify ecosystems most at risk of collapse to prioritise the conservation efforts needed to avert such an event (Keith et al., 2013). Similar to the IUCN Red List of Threatened Species, assessment is against five criteria: A. Reduction in geographic distribution, B. Restricted geographic distribution, C. Environmental degradation, D. Disruption of biotic processes or interactions and E. Quantitative risk analysis. The thresholds have been devised for assessment of ecosystems across their global extent and sub-global assessment guidelines are still under consideration (Bland et al., 2017). However, several national assessments have been undertaken, with the RLE being applied in over 100 countries so far with 4,000 ecosystem units, across various scales, assessed (IUCN Red List of Ecosystems, 2023). In Africa, 920 ecosystems have been assessed across 21 countries (Keith et al., 2023).

There have already been some connections made between the RLE and IPAs. Following IPA identification in the Chiquitanía region of Bolivia, a RLE assessment was undertaken and the threatened habitats within IPAs were retrospectively identified (Martinez-Ugarteche et al., 2023). In the British Virgin Islands, concepts from RLE criteria, including restricted distribution and “continuous decline”, were applied in threatened habitat identification for IPAs (Barrios et al., 2019). Through applying the RLE to identify preliminary IPAs in Uganda, we discuss insights into the applicability of the RLE within IPAs, particularly in the Tropics.

We subsequently assessed the application of the three thresholds of IPA sub-criterion C(iii) at stage two. We consider how application of each of these thresholds might impact which sites trigger this sub-criterion and, therefore, impact the resultant IPA network. This analysis provides insights that can improve the application of criterion C and support the identification of IPA networks that optimally represent threatened habitats. With greater opportunities to conserve these habitats identified, nations can be supported in their commitments towards protecting and restoring ecosystems under the Kunming-Montreal Global Biodiversity Framework.

## Methods

### Study area

Uganda hosts a wide range of habitats, influenced by the country’s large variation in altitude, precipitation, soils and geology (Hamilton, 1984; J. Kalema & Bukenya-Ziraba, 2005). From the formative descriptions of Africa’s phytochoria by White (1983) to more recent updates using statistical methods, such as Linder *et al*. (2005, 2012) and Droissart *et al*. (2018), Uganda has been defined as a transitional zone between major African vegetation types. The country’s vegetation has influences from the Guineo-Congolian, Sudanian, Somalia-Masai and East Afromontane phytochoria (the latter including the Albertine Rift region), each of which extend within Uganda’s borders in their own right.

The most comprehensive account of Uganda’s habitats lists 22 vegetation types (Langdale-Brown et al., 1964). However, some of these habitats are known to have suffered significant losses. For instance, it is estimated that forest and woodland covered around 45% of Uganda’s land area in 1890 (NEMA, 1996). By 1996, the original extent of forest and woodland was estimated to have more than halved to around 20% of Uganda’s area (NEMA, 1996), and these losses continue. Between 2001 and 2022, 1.03 Mha of tree cover (of over 30% canopy) was lost, equating to a 22% decline in area over this period (World Resources Institute, 2023).

The most recent analysis of Uganda’s habitats by Plumptre *et al*. (2021) estimated vegetation loss between 1964 to 2010, using Langdale-Brown *et al*. (1964) as a baseline, and applied RLE criteria to identify nationally threatened habitats. The habitats identified were primarily savanna types, indicating a possible shift from pressure on forest vegetation to savannas in the 20^th^ century.

### Vegetation analyses

Uganda lacks extensive habitat information from before Langdale-Brown et al. (1964). However, we sought to capture loss in habitats, particularly examining reported losses in forest types, that occurred before this time. Without further historical data at a national scale, trends in vegetation were analysed using the “Potential Vegetation for Eastern Africa” (van Breugel et al., 2015), following methods used in Mozambique (Lötter et al., 2023). This map was compared to “The Vegetation of Uganda” as mapped in the 1960s by Langdale-Brown *et al*. (1964) and subsequently the 2017 NFA Land Cover map (NFA, 2017) following the methodology shown in Figure 1.

**Figure 1:**
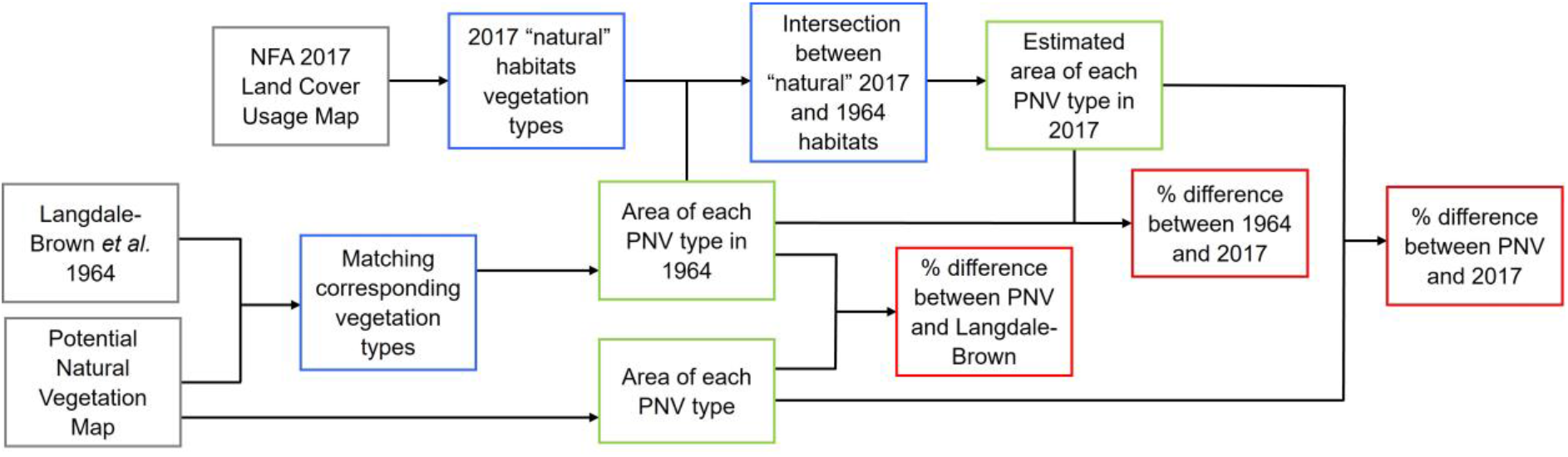
Summary of workflow used to estimate vegetation change pre- and post-1964. Grey boxes indicate data inputs, blue indicate analysis steps, green indicate final GIS layers and red indicate numerical outputs that fed into the Red List of Ecosystems assessment.

To understand the difference between potential natural vegetation (PNV) and the vegetation in 1964, the vegetation categories that represented the same vegetation types between each map were matched (Table S1). Vegetation types identified by Langdale-Brown *et al*. (1964) as anthropogenic were not matched to any PNV type. For instance, *Combretum-Oxytenanthera-Hyparrhenia* savannah is described as likely “derived from Lowland Bamboo Thicket by a combination of bamboo cutting and shifting cultivation” and represents an anthropogenic vegetation type in Uganda that is not assessed here. Through comparing these two maps, we estimated the pre-1960s decline in Uganda’s habitats.

The 2017 NFA Land Use Cover map (NFA, 2017) was used to measure recent vegetation change. This map has higher-level vegetation categories and so could not be used for direct comparison with the PNV and Langdale-Brown *et al*. (1964) maps. However, following the methods of Plumptre *et al*. (2019), the 2017 map was used to identify the extent of “natural” and “anthropogenic” vegetation types across Uganda (Table S2). The extent of each PNV type remaining in the 1960s was subsequently clipped to the extent of the “natural” vegetation types as defined in 2017 by the NFA map, to remove obviously transformed habitats.

GIS analyses were performed using the “*sf”* package in R (Pebesma, 2018).

### Application of Red List of Ecosystems criteria (step one)

Assessments were made according to Bland *et al*. (2017). Vegetation types were assessed against criterion A of the Red List of Ecosystems criteria, “reduction in geographic distribution” (Table 2). Other RLE criteria were not used for assessment as the area-based thresholds under criterion B refer to global-scale assessments that are less relevant nationally, while Criteria C – E require data that is not currently available for Uganda. A1 sub-criterion “Past (over the past 50 years)” was used to assess the percentage change between 1964 and 2017, which is roughly equivalent to 50 years. Following Lötter et al. (2023) we also applied the A3 sub-criterion “Historical (since approximately 1750)” to assess the percentage change between PNV to 2017.

**Table 2:** IUCN Red List of Ecosystems criterion A (Bland *et al*. 2017). Within this analysis sub-criteria A1 and A3 have been applied.

|  | CR | EN | VU |
| --- | --- | --- | --- |
| **A1** Past (over the past 50 years) | ≥ 80% | ≥ 50% | ≥ 30% |
| **A2a** Future (over the next 50 years) | ≥ 80% | ≥ 50% | ≥ 30% |
| **A2b** Any 50 year period (including the past, present and future) | ≥ 80% | ≥ 50% | ≥ 30% |
| **A3** Historical (since approximately 1750) | ≥ 90% | ≥ 70% | ≥ 50% |

The extent of each threatened vegetation type within the protected area network was then measured using the data from the World Database of Protected Areas (UNEP-WCMC & IUCN, 2023).

### Application of IPA criteria and testing of thresholds (step two)

The vegetation types identified as nationally threatened under Red List of Ecosystems criteria were used to identify IPA trigger sites using criterion C(iii).

IPA trigger sites were identified using the estimated 2017 extent of each threatened habitat. All three IPA thresholds (Box 1) were applied to understand how application of each might alter the trigger sites and the resulting IPA network.

While there is deliberately no definition of “best sites”, for the application of five best sites, or “best quality examples”, to collectively prioritise 20% of the national resource to allow flexibility in assessments, “best” and “best quality” was interpreted here as largest continuous, or near-continuous, areas of each vegetation type nationally.

### Delineation of sites

To assess sites against IPA criterion C(iii) and compare thresholds, it was necessary for the purposes of this analysis to standardise site delineation. Vegetation patches of the same type within the same protected area were treated as a single site. If a vegetation patch spanned more than one protected area, then the entire patch was assigned to the protected area which hosted the greatest extent of that vegetation patch. Vegetation patches with less than 10% of their extent within a protected area were not considered “protected” to avoid grouping large expanses of vegetation that are only marginally within a protected area. Finally, any vegetation patch, inside or outside the protected area network, within 500 m of another patch of the same type was clustered together and treated as a single site.

## Results

### Red List of Ecosystems assessment

Eleven vegetation types were found to be nationally threatened with collapse. Lowland bamboo, evergreen and semi-evergreen bushland and thicket and Lake Victoria drier peripheral semi-evergreen Guineo-Congolian rain forest were each found to be Critically Endangered (Table 3). Notably, lowland bamboo declined significantly before and after 1964, meeting the Critically Endangered threshold under both A1 and A3. While the latter two habitat types experienced most declines before 1964, both of these habitats did decline significantly after 1964, meeting the Vulnerable threshold under A1 for this time period.

**Table 3:** Percentage differences in area of each habitat type between potential natural vegetation (PNV), 1960s vegetation and 2017. Red List of Ecosystem assessments were completed based upon “% difference 1964 and 2017” using sub-criterion A1 and on “% difference PNV and 2017” using sub-criterion A3, both of these columns are colour coded according to threat category (red – Critically Endangered, orange – Endangered and yellow – Vulnerable) and the final category is given in the final column.

| Potential Natural Vegetation (PNV) |  |  |  | % | % | % | RLE |
| --- | --- | --- | --- | --- | --- | --- | --- |
|  |  |  |  | difference | difference | difference | category |
|  | PNV area<br>(km<sup>2</sup>) | 1964 area<br>(km<sup>2</sup>) | 2017 area<br>(km<sup>2</sup>) | PNV and<br>1964 | 1964 and<br>2017 (A1) | PNV and<br>2017 (A3) |  |
| 1. Lowland bamboo | 1,406.14 | 300.21 | 52.61 | -78.65% | -82.48% | -96.26% | CR A1+3 |
| 2. Evergreen and semi-evergreen bushland and thicket | 20,637.03 | 1,616.64 | 856.19 | -92.17% | -47.04% | -95.85% | CR A3 |
| 3. Lake Victoria drier peripheral semi-evergreen<br>Guineo-Congolian rain forest | 40,268.71 | 6,611.49 | 3,881.93 | -83.58% | -41.29% | -90.36% | CR A3 |
| 4. Moist *Combretum* wooded grassland | 17,603.28 | 15,668.64 | 3,169.73 | -10.99% | -79.77% | -81.99% | EN A1+3 |
| 5. Afromontane rain forest | 4,381.01 | 1,706.22 | 1,000.54 | -61.05% | -41.36% | -77.16% | EN A3 |
| 6. Palm wooded grassland | 2,758.90 | 2,484.28 | 1,011.47 | -9.95% | -59.29% | -63.34% | EN A1 |
| 7. *Vitellaria (Butyrospermum)* wooded grassland | 25,059.42 | 22,785.89 | 7,332.85 | -9.07% | -67.82% | -70.74% | EN A1 |
| 8. Afromontane undifferentiated forest | 2,791.26 | 1,399.13 | 1,285.82 | -49.87% | -8.10% | -53.93% | VU A3 |
| 9. Dry *Combretum* wooded grassland | 43,024.15 | 41,499.72 | 21,540.83 | -3.54% | -48.09% | -49.93% | VU A1+3 |
| 10. *Vitex-Phyllanthus-Shirakipsis (Sapium)-Terminalia*<br>*and Terminalia glaucescens* woodland | 3,394.34 | 3,376.88 | 1,869.27 | -0.51% | -44.65% | -44.93% | VU A1 |
| 11. Freshwater swamp | 8,513.43 | 7,112.47 | 4,706.56 | -16.46% | -33.83% | -44.72% | VU A1 |
| 12. Upland *Acacia* wooded grassland | 626.38 | 462.09 | 373.84 | -26.23% | -19.10% | -40.32% | LC |
| 13. Somalia-Masai *Acacia-Commiphora* deciduous bushland and thicket | 8,261.81 | 6,260.91 | 5,200.60 | -24.22% | -16.94% | -37.05% | LC |
| 14. Riverine wooded vegetation | 680.93 | 680.73 | 505.71 | -0.03% | -25.71% | -25.73% | LC |
| 15. Edaphic grassland and wooded grassland on drainage-impeded or seasonally flooded soils | 23,570.14 | 24,015.23 | 17,302.93 | 1.89% | -27.95% | -26.59% | LC/NT A1 |
| 16. Swamp forest | 274.71 | 278.96 | 249.95 | 1.55% | -10.40% | -9.01% | LC |
| 17. Montane Ericaceous belt | 418.93 | 386.53 | 382.09 | -7.73% | -1.15% | -8.79% | LC |
| 18. Climatic grasslands | 797.79 | 797.64 | 794.42 | -0.02% | -0.40% | -0.42% | LC |
| 19. Afroalpine vegetation | 273.57 | 288.50 | 278.91 | 5.46% | -3.32% | 1.95% | LC |
| 20. Afromontane Bamboo and *Hagenia* | 533.29 | 575.85 | 548.65 | 7.98% | -4.72% | 2.88% | LC |

Four vegetation types were found to meet the Endangered threshold: moist *Combretum* wooded grassland, Afromontane rain forest, palm wooded grassland and *Vitellaria (Butyrospermum)* wooded grassland. While the majority of decline in Afromontane rainforest occurred pre-1964, for the other Endangered habitats, all wooded grassland (savanna) types, almost all of their decline occurred post-1964. Similarly, of the five Vulnerable habitats the only forest type, Afromontane undifferentiated forest, the only type to show most of its decline before 1964. Uniquely among all the threatened vegetation types, Afromontane undifferentiated forest has remained relatively stable since 1964, showing only 8.1% decline (equivalent to 113 km^2^ lost).

### Distribution and protection of threatened vegetation types

Figure 2a shows the distribution, according to the 2017 estimate, of each of the vegetation types threatened with collapse, while 2b illustrates the potential extent of these vegetation types. In total, threatened vegetation types cover 16.78% of Uganda’s area, and 20.21% of the country’s terrestrial area. Contrastingly, the potential extent of these threatened vegetation types covers 65.63% of the total country area, and 78.93% of the terrestrial area.

**Figure 2:**
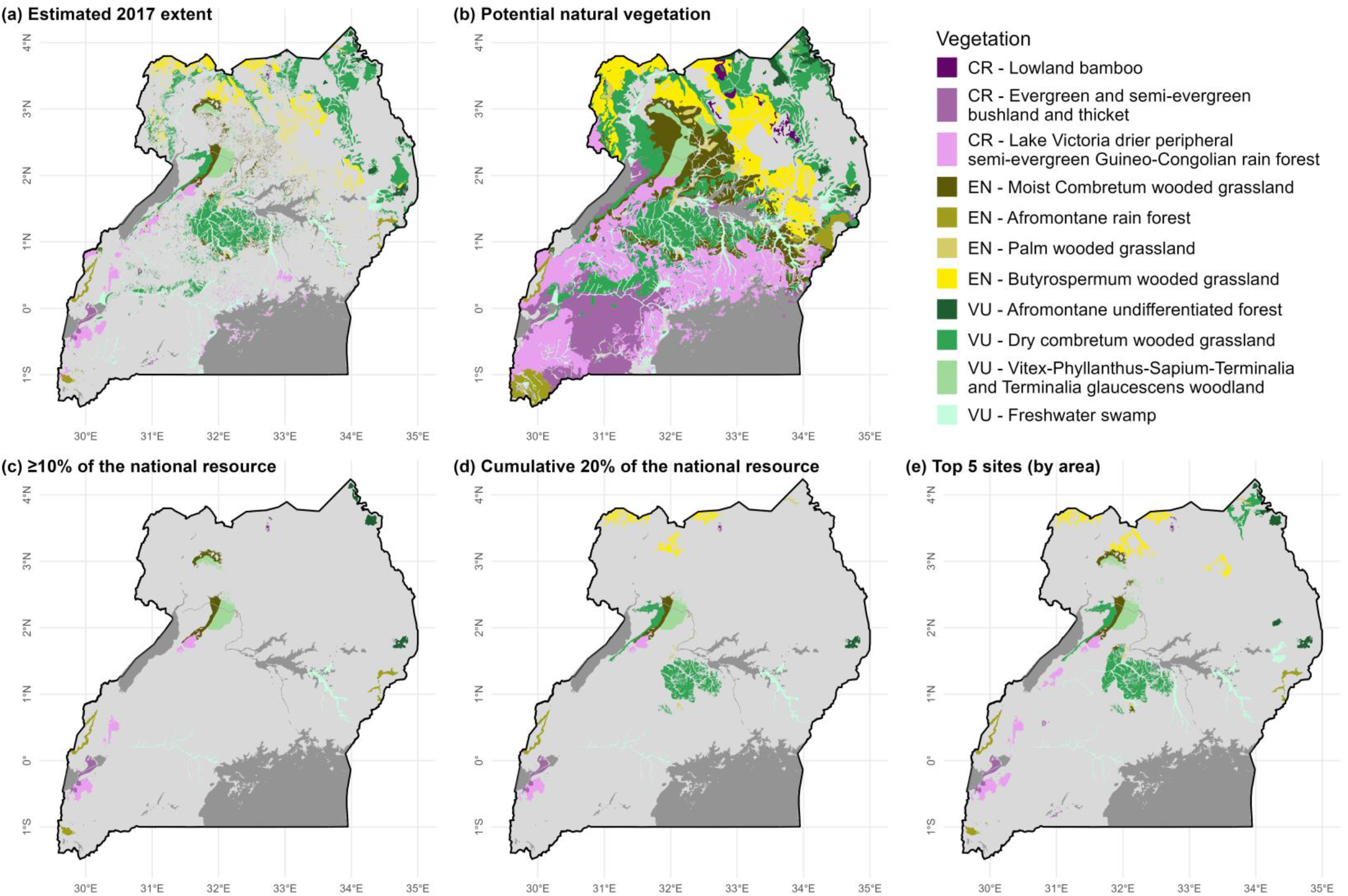
The distribution of the threatened habitats of Uganda. (a) The estimated 2017 extent of each threatened habitat; (b) potential natural extent of each threatened habitat; (c – e) the sites identified to trigger IPAs based on each criterion C(iii) threshold.

**Figure 3:**
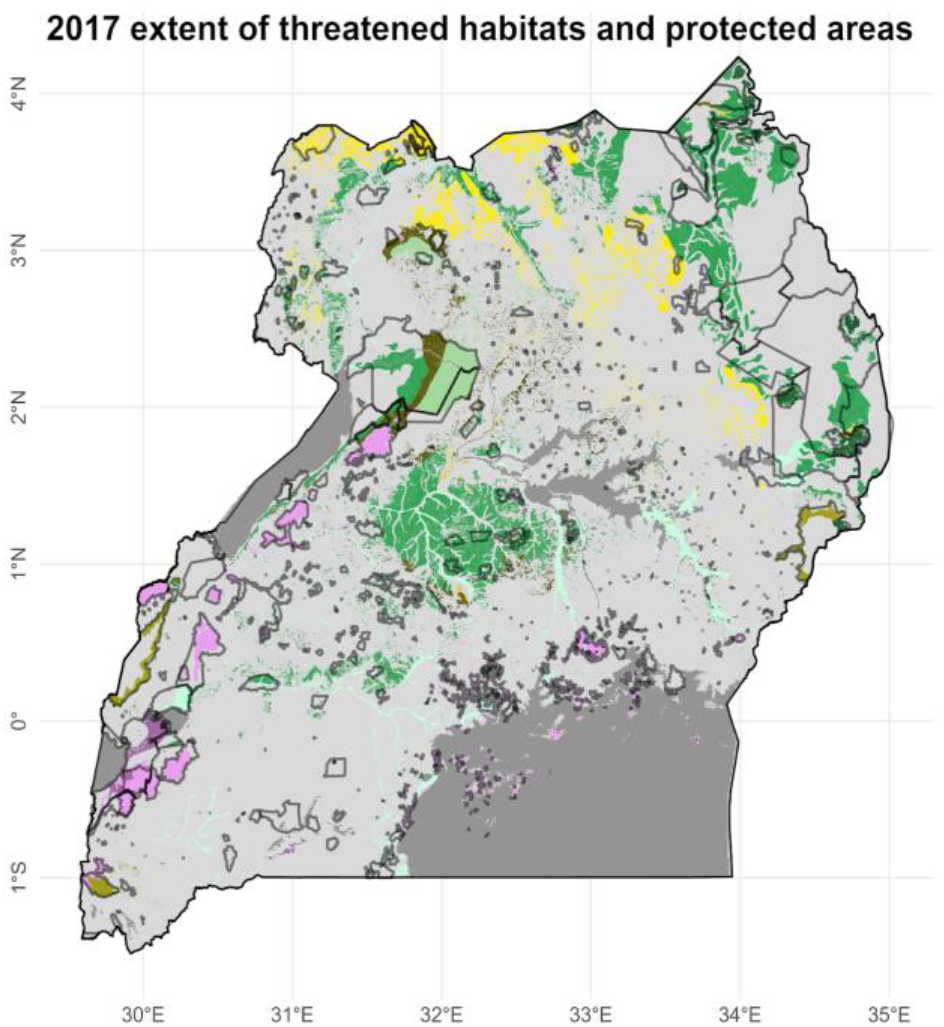
2017 extent of threatened habitats of Uganda mapped alongside the protected area network (grey outline). Colour legend to habitats in Figure 2.

36.78% of the 2017 estimate of the extent of all threatened vegetation types fall within protected areas; Table 4 has a breakdown of protected area coverage by vegetation type. The 2017 extent of some vegetation types, particularly forest types, is well represented within the protected area network. However, both Afromontane rainforest and Lake Victoria drier peripheral semi-evergreen Guineo-Congolian rainforest have much larger potential extents outside of the current protected area network. Lowland bamboo, the most threatened vegetation according to the RLE analysis, has the smallest percentage extent within the protected area network. Interestingly, is the only vegetation type to have a larger potential extent within protected areas than is estimated for 2017. This reflects several forest reserves in Karamoja where bamboo is thought to potentially occur, but were mapped as dry *Combretum* savanna, derived from lowland bamboo by Langdale-Brown (1964).

**Table 4:** The proportion of each vegetation type within protected areas.

| **Vegetation type** | **% of 2017 area**<br>**within protected**<br>**areas** | **% of PNV area**<br>**within protected**<br>**areas** |
| --- | --- | --- |
| EN - Afromontane rainforest | 91.69% | 25.27% |
| VU - Afromontane undifferentiated forest | 80.58% | 74.62% |
| CR - Lake Victoria drier peripheral semi-evergreen<br>Guineo-Congolian rain forest | 79.81% | 14.66% |
| VU - *Vitex-Phyllanthus-Shirakiopsis (Sapium)-Terminalia*<br>and *Terminalia glaucescens* woodland | 68.69% | 41.45% |
| CR - Evergreen and semi-evergreen bushland and thicket | 66.57% | 12.24% |
| EN - Moist *Combretum* wooded grassland | 30.00% | 7.57% |
| VU - Freshwater swamp | 20.08% | 12.87% |
| VU - Dry *Combretum* wooded grassland | 19.79% | 16.75% |
| EN - Palm wooded grassland | 8.50% | 5.06% |
| EN - *Vitellaria (Butyrospermum)* wooded grassland | 7.25% | 4.08% |
| CR - Lowland bamboo | 4.31% | 33.72% |

### Application of IPA thresholds

When testing the IPA sub-criterion C(iii) thresholds, our analysis finds that the application of the quantitative thresholds (site contains ≥10% of the national resource or site is among the best quality examples required to collectively prioritise up to 20% of the national resource) yields fewer sites than application of the qualitative “five best sites” threshold (Table 5). Vegetation types with many small sites are not selected through the ≥10% threshold, as is the case for palm wooded grassland, while vegetation types with a small number of sites that account for a large proportion of the national resource might have few sites qualifying under either the ≥10% or cumulative 20% thresholds. Evergreen and semi-evergreen bushland and thicket, for instance, has a single site accounting for 58% of the national resource, while all others represent less than 10%, so both quantitative criteria only trigger a single site. Across all threatened habitats, even with both quantitative thresholds combined, this does not, apart from in the case of palm wooded grassland, result in five sites being selected.

**Table 5:** Number of IPAs triggered by the three different thresholds of Criterion C. The use of the quantitative thresholds, “≥10% of national resource” and “best quality examples required to collectively prioritise up to 20% of the national resource”, leads to fewer sites being identified compared to the qualitative threshold of “5 best sites”.

| Vegetation type | ≥10% of<br>national<br>resource | Cml. 20% of the<br>national<br>resource | Total sites<br>captured by<br>quantitative<br>thresholds | The 5 best sites<br>nationally |
| --- | --- | --- | --- | --- |
| CR – Lowland bamboo | 2 | 1 | 2 | 5 |
| CR – Evergreen and semi-evergreen<br>bushland and thicket | 1 | 1 | 1 | 5 |
| CR – Lake Victoria drier peripheral<br>semi-evergreen Guineo-Congolian<br>rain forest | 3 | 2 | 3 | 5 |
| EN – Moist *Combretum* wooded<br>grassland | 2 | 1 | 2 | 5 |
| EN – Afromontane rain forest | 3 | 1 | 3 | 5 |
| EN – Palm wooded grassland | 0 | 5 | 5 | 5 |
| EN – *Vitellaria (Butyrospermum)*<br>wooded grassland | 0 | 3 | 3 | 5 |
| VU – Afromontane undifferentiated<br>forest | 3 | 1 | 3 | 5 |
| VU – Dry *Combretum* wooded<br>grassland | 0 | 3 | 3 | 5 |
| VU – *Vitex-Phyllanthus-Shirakiopsis*<br>(*Sapium*)-*Terminalia* and *Terminalia*<br>*glaucescens* woodland | 2 | 1 | 2 | 5 |
| VU – Freshwater swamp | 2 | 2 | 2 | 5 |

## Discussion

### Threatened habitats of Uganda

Uganda has experienced severe declines in its natural vegetation types, with over half now threatened with collapse. Figures 2a and 2b demonstrate how vegetation types that may have once dominated Uganda have declined to small fraction of the nation’s land area. Clearance of land for agriculture, demand for charcoal, fuelwood and timber, and expansion of urban and industrial centres are all currently linked to the conversion of habitats in Uganda (NEMA, 2016).

Lowland bamboo vegetation is the closest to collapse with an estimated 52.61 km^2^ of this habitat remaining. While the bamboo species that forms these stands is not rare - *Oxytenanthera abyssinica* is native throughout tropical Africa - this habitat type is very rare nationally, with even the predicted extent very limited in area when compared to other types. Moreover, this habitat type is estimated to be rare within the East Africa floral region, with only Uganda hosting any areas of this habitat (van Breugel et al., 2015). One of the key purposes of Criterion C of the IPA criteria is to capture threatened and extremely restricted habitats “regardless of how botanically rich they are” (Darbyshire et al., 2017). The example of lowland bamboo habitat in Uganda demonstrates how application of this criterion is an important opportunity to capture botanically rare areas that are otherwise poor in rare or threatened species required to trigger Criteria A and B.

The habitats identified as threatened are otherwise largely forest and wooded grassland types. Forest clearance in Uganda has been well-documented. The earliest evidence of forest clearance in Uganda originated from at least 2,000 years ago (Chapman & Chapman, 1996; Hamilton, 1984). In more recent history, the British Protectorate promoted the clearance of forests for introduced cash crops such as coffee, tea and sugar from the 1890s, while valuable hardwood stands were designated Crown Forests (Turyahabwe & Banana, 2008). These forests were managed under a forestry regime which at times included supplementary planting of desirable species and arboricide use on less commercially viable trees (Hamilton, 1984; Howard, 1991). The impact of the latter appears not to have been documented within Langdale-Brown *et al*. (1964), and as such loss of natural forest stands may well have been underestimated within this analysis.

It is notable that forest decline largely occurred before 1964 (Table 1). By the 1960s, much of the remaining forest resource was already within protected areas (Langdale-Brown et al., 1964; UNEP-WCMC & IUCN, 2023). This analysis appears to demonstrate that the establishment of these reserves has been reasonably successful in securing forest resources, accounting for much of the extent of forest habitats today (Figure 2). However, many of the forest remnants outside protected areas, and inside more poorly managed protected areas, have been lost since 1964. Even forest in protected areas thought to be secure can quickly become at risk. For example, since the 2017 land use cover map was produced, large areas of Bugoma Central Forest Reserve have been lost to sugar plantation (Tenywa, 2021). This site is a key area for the Critically Endangered Lake Victoria drier peripheral semi-evergreen Guineo-Congolian rain forest and triggers IPA criterion C(iii) as one of the five best sites nationally for this habitat. Unfortunately, this is not an incidence unique to Uganda and the pressure for economic development in many tropical nations puts pressure on forests that results in protection in name only (Bongben, 2023; Curran et al., 2004; Graham, 2022). Also of concern are the threatened wooded grassland (savanna) habitat types that are mostly outside protected areas. As Uganda’s protected area network was being established, the key priority of the British Protectorate was the management of forest resources and regulation of game hunting (Banana et al., 2018). The designation of game reserves did lead to some threatened savanna vegetation being designated within protected areas, including the Murchison Falls National Parks (originally designated a game reserve) which host significant areas of both moist (EN) and dry (VU) *Combretum* savanna and *Vitex-Phyllanthus-Shirakiopsis-Terminalia* and *Terminalia glaucescens* woodland (VU). Tropical savannas and dry forests are globally known to be underrepresented within protected area networks (Pennington et al., 2018). If the conservation value of these vegetation types was not considered independently of other aspects of biodiversity, this may well explain the generally lower levels of protected area coverage for savanna vegetation types in Uganda.

The post-1960s decline in savanna habitats supports the assessment by Plumptre et al. (2021) for the last 50 years which finds that threatened habitats consist exclusively of savanna grassland and woodland habitats. These two studies also substantiate anecdotal evidence of greater clearance for agriculture and pastureland in more recent decades within these habitats (Banana et al., 2014; NEMA, 2016).

With this growing evidence, there is a need for greater recognition of the threats to savanna habitats. These habitats are also very important in provision of ecosystem services for Ugandan communities including provision of fuel, construction materials, medicines and food (V. N. Kalema, 2010). Across sub-Saharan Africa, high rainfall savannas are expected to represent the next frontier for commercial agriculture expansions (Pennington et al., 2018), and so we might expect some declines to continue unless action is taken to conserve them. A greater focus on these threatened habitats, through prioritisation schemes such as IPAs, can, however, highlight the need for conservation of such ecosystems.

### Application of the Red List of Ecosystems

A limitation of the Red List of Ecosystems criteria is the need for data that may not be available in the Tropics. In table 1 we see how many tropical nations lack the vegetation data or expertise required to identify threatened habitats systematically for IPAs. While there has been progress in assessing ecosystems in 21 African nations, there are still significant gaps (Keith et al., 2023).

The use of potential natural vegetation has allowed us to identify trends in habitat loss before the first comprehensive habitat maps of Uganda were produced in the mid-20th century. This methodology follows Lötter et al. (2023) in applying RLE sub-criterion A3 to PNV data as the most appropriate threshold with which to interpret the data. IUCN incorporated A3 into the RLE criteria to recognise the negative impact of historical declines on present day resilience of an ecosystem (Bland et al., 2015). Although it is widely accepted that there is a longstanding history of human influence on the vegetation of Uganda (Chapman & Chapman, 1996; Hamilton, 1984), past declines in habitat types will have negatively impacted their resilience in the present and they should therefore be recognised as threatened habitats on this basis.

While Uganda, alongside several other nations in eastern Africa, is fortunate to have a PNV map, several other countries within the tropics do not have this data. However, though the lack of historical vegetation maps cannot be redressed, PNV maps could be developed for other countries in the Tropics to allow for the long-term changes in vegetation to be examined through RLE assessments. This was the approach taken in Mozambique, where a PNV map was developed for RLE assessment by combining the limited historical vegetation data with species distribution data, remote sensing and other environmental data using both supervised learning algorithms and expert-input (Lötter et al., 2023). Application of PNV data could allow for greater understanding of trends in vegetation in the Tropics and, if deemed appropriate, IUCN could consider formal integration of PNV data with bespoke thresholds or guidance.

### IPA identification

Using the RLE, we have identified habitats that are likely to have declined severely and are threatened with collapse. IPAs can play an important role in identifying priority sites at which to conserve such habitats.

Other prioritisation schemes, such as Important Bird Areas, do not incorporate habitat-specific criteria, while Key Biodiversity Areas do have the criteria to include ecosystems but rely on global Red List of Ecosystem assessments (of global ecosystems or, at a broader scale, biogeographic ecotypes) that are yet to be completed (KBA, 2023; KBA Standards and Appeals Committee, 2020). IPAs, in contrast, have the flexibility to identify threatened and restricted habitats on a regional and national scale. This inclusion of sub-global habitat types has enabled conservation priorities to be identified and integrated into IPA networks across several countries over the past two decades which, in turn, has informed policymakers and other conservation stakeholders. In Guinea, for example, identification of threatened habitats within IPAs has recently led to their integration within conservation action plans for two IPAs, Diécké Classified Forest, and Mount Bero Classified Forest, Nzérékoré and Beyla (Couch, 2022; Woreto et al., 2022).

Despite successes, the application of habitat thresholds under IPA criterion C has not yet been tested. As demonstrated in the Table 1, different IPA projects have applied criterion C differently, largely due to differences in perceived or actual data availability, expertise available, and stakeholder engagement. However, it has previously not been examined how this differing application could change outcomes in the IPA networks themselves.

We demonstrate here that application of the quantitative thresholds only, limits the number of IPA trigger sites that can be identified. Using the “≥10% of the national resource” threshold biases against heavily fragmented but relatively widespread vegetation types, such as palm wooded grassland in Uganda, where there are no sites that account for large proportions of the national resource. While application of the “20% cumulative” threshold as well as the “≥10% national resource” threshold would overcome this particular issue, both of these thresholds produce sub-optimal outcomes for habitats that are highly restricted and have one particularly large site that alone accounts for ≥20% of the national resource. This is the case with evergreen and semi-evergreen bushland and thicket with a single large site in Queen Elizabeth National Park that accounts for 58% of the national resource. While Queen Elizabeth National Park is the most feasible opportunity to conserve this vegetation type nationally, the application of quantitative thresholds alone prevents consideration of other sites, possibly including those where restoration efforts could see this habitat expand.

The IPA criteria, however, state that “the selection of best sites should only be applied where quantitative data are not available and cannot be inferred” (Darbyshire et al., 2017). We have demonstrated here how this guidance effectively limits the number of IPA trigger sites when quantitative data are available. We instead suggest that the “five best sites” threshold is available to use for both qualitative and quantitative data. Some IPA programmes have already implemented this approach, including the British Virgin Islands and the Mozambique IPA projects (Barrios et al., 2019; Darbyshire et al., 2023). Percentage national resource should, however, still be reported when using the five best sites threshold for transparency, and to enable practitioners to plan and prioritise conservation actions within and between sites. Even where an IPA does not qualify under criterion C, but meets other criteria, reporting any threatened habitats present within the site would be informative.

### Future steps

In this study, we have preliminarily identified the threatened habitats and IPAs for Uganda. However, it is important to seek the expertise and views of stakeholders, consider other IPA criteria met and any complementarity between sites when identifying IPAs. In addition, ground-truthing or more localised data could validate the threat status of a vegetation type as well as identifying which sites are best to conserve these habitats.

In this study, further research is particularly needed for lowland bamboo habitat where the areas are so small and fragmented that they can scarcely be estimated accurately without site surveys. The finding that there is much greater potential for this habitat to occur in protected areas than has been realised (Table 4), is particularly interesting and may suggest either an inaccuracy in estimated extent of habitat or that management interventions are required to promote the presence of this habitat within protected areas. Contrasting with the highly limited lowland bamboo habitat, we note that there are extensive areas of palm wooded grassland within Murchison Falls National Park which are not accounted for within the vegetation maps used (J. Kalema, pers. obs.). Further consideration, including consultation of experts and additional evidence, will therefore be undertaken to incorporate threatened habitats within IPAs in Uganda.

## Conclusions

Uganda has committed, through its National Biodiversity Strategy Action Plan II 2015 – 2025 to “identify, map and prioritize degraded habitats including forests and wetlands” to “restore degraded natural habitats” (NEMA, 2016). Commitment to the Kunming-Montreal Global Biodiversity Framework further requires Uganda, and all signatories globally, to enhance and expand the area of natural ecosystems by 2050 and bring 30% of degraded ecosystems into restoration by 2030. Improving the recognition of threatened habitats within IPAs is therefore crucial to ensuring maximal contribution towards post-2020 conservation goals. While there are some challenges in applying the RLE within IPAs, PNV may offer a solution to limited data availability when identifying threatened habitats. Importantly, this study finds that the exclusion of the “five best sites” threshold when assessing IPAs with quantitative habitat data is limiting conservation opportunities. We instead recommend that this threshold should be available for use with both qualitative and quantitative habitat data to support practitioners in maximising the inclusion of threatened habitats within IPAs.

## Author Contribution Statement

SLR designed the study. SLR designed the methodology which was further developed with contributions from ID, JW and JK. Analysis and data visualisation was performed by SLR. The first draft of the manuscript was drafted by SLR and SO, ID and JK commented on previous versions of the manuscript. All authors read and approved the near-final manuscript.

## Target audience

This research will support botanical conservation researchers in the application of IPAs globally and is informative for conservation practitioners in Uganda.

## Acknowledgements

The Tropical Important Plant Areas of Uganda project was generously supported by the players of the People’s Postcode Lottery, the Woodspring Trust and the P.F. Charitable Trust.

## Data availability statement

We will deposit GIS layers created in this study as shapefiles.

## Notes

### Competing Interest Statement

The authors have declared no competing interest.

## Bibliography

Anderson, C. (2002). *Identifying Important Plant Areas*. www.plantlife.org.uk

Banana, A. Y., Byakagaba, P., Russell, A. J. M., Waiswa, D., & Bomuhangi, A. (2014). A review of Uganda’s national policies relevant to climate change adaptation and mitigation: Insights from Mount Elgon. In *Centre for International Forestry Research*. https://www.jstor.org/stable/resrep02362.7

Banana, A. Y., Nsita, S., & Bomuhangi, A. (2018). Histories and genealogies of Ugandan forest and wildlife conservation; the birth of the protected area estate. In Sandbrook Chris, Connor Joseph Cavanagh, & David Mwesigye Tumusiime (Eds.), Conservation and development in Uganda. Routledge.

Barrios, S., Clubbe, C., Dani Sanchez, M., Grant, K., Hamilton, M. A., Harrigan, N., Heller, T., Abbott, J. S., Smith, T., Varlack, L., Pascoe, N. W., Martin, D., & Hamilton, A. (2019). *Identifying and Conserving Tropical Important Plant Areas in the British Virgin Islands (2016-2019): Final Technical Report*.

Bland, L. M., Collen, B., Orme, C. D. L., & Bielby, J. (2015). Predicting the conservation status of data-deficient species. Conservation Biology, 29(1), 250–259. 10.1111/cobi.12372

Bland, L. M., Keith, D. A., Miller, R. M., Murray, N. J., & Rodríguez, J. P. (2017). *Guidelines for the application of IUCN Red List of Ecosystems Categories and Criteria Edited by*. 10.2305/IUCN.CH.2016.RLE.3.en

Bongben, L. (2023, August 1). Cameroon government again opens way for logging in Ebo Forest. Mongabay. https://news.mongabay.com/2023/08/cameroon-government-again-opens-way-for-logging-in-ebo-forest/

CBD. (2022). *Kunming-Montreal Global Biodiversity Framework. CBD/COP/DEC/15/4*. https://www.cbd.int/doc/decisions/cop-15/cop-15-dec-04-en.pdf

Chapman, C. A., & Chapman, L. J. (1996). Mid-elevation Forests: A history of Disturbance and Regeneration. In T. R. McClanahan & T. P. Young (Eds.), East African Ecosystems and Their Conservation. Oxford University Press.

Couch, C. (2022). *Plantes de la Foret Classee de Mt Bero, Prefectures de Nzerekore et Beyla. Produit du Projet “Elargissement des Aires Protégées en Guinée y Compris les Zones Tropicales Importantes pour les Plantes (ZTIPS)*. https://kew.iro.bl.uk/concern/reports/fbd8ad02-a6a6-408e-ba03-ddc121093ad6

Couch, C., Cheek, M., Haba, P., Molmou, D., Williams, J., Magassouba, S., Doumbouya, S., & Diallo, M. Y. (2019). Threatened Habitats and Tropical Important Plant Areas (TIPAs) of Guinea, West Africa. 10.34885/169

Curran, L. M., Trigg, S. N., McDonald, A. K., Astiani, D., Hardiono, Y. M., Siregar, P., Caniago, I., & Kasischke, E. (2004). Lowland Forest Loss in Protected Areas of Indonesian Borneo. Science, 303(5660), 1000–1003. 10.1126/SCIENCE.1091714/SUPPL_FILE/CURRAN.SOM.PDF

Darbyshire, I., Anderson, S., Asatryan, A., Byfield, A., Cheek, M., Clubbe, C., Ghrabi, Z., Harris, T., Heatubun, C. D., Kalema, J., Magassouba, S., McCarthy, B., Milliken, W., de Montmollin, B., Lughadha, E. N., Onana, J. M., Saïdou, D., Sârbu, A., Shrestha, K., & Radford, E. A. (2017). Important Plant Areas: revised selection criteria for a global approach to plant conservation. In Biodiversity and Conservation (Vol. 26, Issue 8, pp. 1767–1800). Springer Netherlands. 10.1007/s10531-017-1336-6

Darbyshire, I., Richards, S., Osborne, J., Matimele, H., Langa, C., Datizua, C., Massingue, A., Rokni, S., Williams, J., Alvez, T., & de Sousa, C. (2023). The Important Plant Areas of Mozambique. Royal Botanic Gardens, Kew.

Droissart, V., Dauby, G., Hardy, O. J., Deblauwe, V., Harris, D. J., Janssens, S., Mackinder, B. A., Blach-Overgaard, A., Sonké, B., Sosef, M. S. M., Stévart, T., Svenning, J.-C., Wieringa, J. J., & Couvreur, T. L. P. (2018). Beyond trees: Biogeographical regionalization of tropical Africa. Journal of Biogeography, 45(5), 1153–1167. 10.1111/jbi.13190

Graham, T. (2022, October 19). Bolivian gold miners push into national park despite country’s green rhetoric. The Guardian. https://www.theguardian.com/world/2022/oct/19/bolivia-gold-miners-amazon-madidi

Hamilton, A. C. (1984). Deforestation in Uganda. Oxford University Press.

House, E., Darbyshire, I., Lukeal, E., Nemomissa, S., Awas, T., Belay Telake, B., & Demissew, S. (2023). *Tropical Important Plant Areas Explorer: Ethiopia*. https://tipas.kew.org/

Howard, P. (1991). Nature Conservation in Uganda’s Tropical Forest Reserves. IUCN, Gland, Switzerland and Cambridge, UK.

IPBES. (2019). Summary for policymakers of the global assessment report on biodiversity and ecosystem services. 10.5281/ZENODO.3553579

IUCN Red List of Ecosystems. (2023). *IUCN Red List of Ecosystems Database*. https://assessments.iucnrle.org/

Kalema, J., & Bukenya-Ziraba, R. (2005). Patterns of plant diversity in Uganda. Biologiske Skrifter, 55, 331–341.

Kalema, V. N. (2010). *DIVERSITY, USE AND RESILIENCE OF WOODY SPECIES IN A MULTIPLE LAND USE EQUATORIAL AFRICAN SAVANNA, CENTRAL UGANDA*.

KBA. (2023). *Key Biodiversity Areas Global Dataset*. https://www.keybiodiversityareas.org/kba-data

KBA Standards and Appeals Committee. (2020). Guidelines for using A global standard for the identification of Key Biodiversity Areas (KBA Standards and Appeals Committee of the IUCN Species Survival Commission and IUCN World Commission on Protected Areas, Ed.; Version 1.1). IUCN. 10.2305/IUCN.CH.2020.KBA.1.1.en

Keith, D. A., Ghoraba, S. M. M., Kaly, E., Jones, K. R., Oosthuizen, A., Obura, D., Costa, H. M., Daniels, F., Duarte, E., Grantham, H., Gudka, M., Norman, J., Shannon, L. J., Skowno, A., & Ferrer-Paris, J. R. (2023). Contributions of Red Lists of Ecosystems to risk-based design and management of protected and conserved areas in Africa. Conservation Biology. 10.1111/COBI.14169

Keith, D. A., Rodríguez, J. P., Rodríguez-Clark, K. M., Nicholson, E., Aapala, K., Alonso, A., Asmussen, M., Bachman, S., Basset, A., Barrow, E. G., Benson, J. S., Bishop, M. J., Bonifacio, R., Brooks, T. M., Burgman, M. A., Comer, P., Comín, F. A., Essl, F., Faber-Langendoen, D., … Zambrano-Martínez, S. (2013). Scientific Foundations for an IUCN Red List of Ecosystems. PLOS ONE, 8(5), e62111. 10.1371/JOURNAL.PONE.0062111

Langdale-Brown, I., Osmaston, H. A., & Wilson, J. G. (1964). The Vegetation of Uganda and its Bearing on Land-Use. Government of Uganda.

Linder, H. P., de Klerk, H. M., Born, J., Burgess, N. D., Fjeldså, J., & Rahbek, C. (2012). The partitioning of Africa: statistically defined biogeographical regions in sub-Saharan Africa. Journal of Biogeography, 39(7), 1189–1205. 10.1111/J.1365-2699.2012.02728.X

Linder, H. P., Lovett, J., Mutke, J. M., Barthlott, W., Jürgens, N., Rebelo, T., & Küper, W. (2005). A numerical re-evaluation of the sub-Saharan phytochoria of mainland Africa. Biologiske Skrifter, 55, 229–252.

Lötter, M., Burrows, J., Jones, K., Duarte, E., Costa, H., McCleland, W., Stalmans, M., Schmidt, E., Darbyshire, I., Richards, S., Soares, M., Grantham, H., Matimele, H., Sousa, C., Alves, T., Zolho, R., Nicolau, D., Ribeiro, N., Macamo, C., … Bandeira S. (2023). *Historical vegetation map and red list of ecosystems assessment for Mozambique – Version 2.0 – Final report*.

Martinez-Ugarteche, M. T., Villarroel, D., Toledo, M., Michme, G., & Klitgaard, B. (2023). Threatened and priority habitats for conservation in the Chiquitano Dry Forest ecoregion, Santa Cruz, Bolivia. Kempffiana, 19(2), 17–68.

Murphy, B., Onana, J. M., van der Burgt, X. M., Ngansop Tchatchouang, E., Williams, J., Tchiengué, B., & Cheek, M. (2023). Important Plant Areas of Cameroon. Royal Botanic Gardens, Kew.

NEMA. (1996). *State of the Environment Report for Uganda*.

NEMA. (2016). *National Biodiversity Strategy and Action Plan II*. http://www.nemaug.org NFA. (2017). *Land Cover Trends in Uganda*.

Pebesma, E. (2018). Simple Features for R: Standardized Support for Spatial Vector Data. The R Journal, 10(1), 439–446. 10.32614/RJ-2018-009

Pennington, R. T., Lehmann, C. E. R., & Rowland, L. M. (2018). Tropical savannas and dry forests. Current Biology, 28(9), R541–R545. 10.1016/J.CUB.2018.03.014

Plumptre, A. J., Ayebare, S., Behangana, M., Forrest, T. G., Hatanga, P., Kabuye, C., Kirunda, B., Kityo, R., Mugabe, H., Namaganda, M., Nampindo, S., Nangendo, G., Nkuutu, D. N., Pomeroy, D., Tushabe, H., & Prinsloo, S. (2019). Conservation of vertebrates and plants in Uganda: Identifying Key Biodiversity Areas and other sites of national importance. Conservation Science and Practice, 1(2). 10.1111/CSP2.7

Plumptre, A. J., Ayebare, S., Kujirakwinja, D., & Segan, D. (2021). Conservation planning for Africa’s Albertine Rift: conserving a biodiverse region in the face of multiple threats. Oryx, 55(2), 302–310. 10.1017/S0030605319000218

Shaltout, K., & Eid, E. (2016). *Important Plant Areas in Egypt with Emphasis on the Mediterranean Region* (Issue April). https://www.researchgate.net/publication/314231880_Important_Plant_Areas_in_Egypt_With_Emphasis_on_the_Mediterranean_Region

Tenywa, G. (2021, May 5). Bitter-sweet exchange: forest cleared for sugarcane - Part 1 - New Vision Official. New Vision. https://www.newvision.co.ug/category/report/bitter-sweet-exchange-forest-cleared-for-suga-NV_101150

Turyahabwe, N., & Banana, A. Y. (2008). An overview of history and development of forest policy and legislation in Uganda. Source: The International Forestry Review, 10(4), 641–656. https://about.jstor.org/terms

UNEP-WCMC, & IUCN. (2023). *World Database of Protected Areas (2023)*. https://www.protectedplanet.net/en/thematic-areas/wdpa

van Breugel, P., Kindt, R., Lillesø, J., Bingham, M., Demissew, S., Dudley, C., Friis, I., Gachathi, F., Kalema, J., Mbago, F., Moshi, H., Mulumba, J., Namaganda, M., Ndangalasi, H., Ruffo, C., Védaste, M., Jamnadass, R., & Graudal, L. (2015). *Potential Natural Vegetation Map of Eastern Africa (Burundi, Ethiopia, Kenya, Malawi, Rwanda, Tanzania, Uganda and Zambia). Version 2.0*. Forest & Landscape Denmark and World Agroforestry Centre (ICRAF). http://vegetationmap4africa.org

White, F. (1983). The vegetation of Africa. A descriptive memoir to accompany the Unesco / AETFAT / UNSO vegetation map of Africa. 10.5281/zenodo.293797

Woreto, D., Couch, C., & Simbiano, F. J. (2022). *Plan d’action de Conservation Pour les Plantes de la Foret Classee de Diecke, Prefecture de Yomou. Produit du Projet “Elargissement des Aires Protégées en Guinée y Compris les Zones Tropicales Importantes pour les Plantes (ZTIPS).* https://kew.iro.bl.uk/concern/reports/cc6e584f-a98c-4ece-b2be-617dfbc62bee

World Resources Institute. (2023). *Global Forest Watch*. https://www.globalforestwatch.org/

Yahi, N., Vela, E., Benhouhou, S., Belair, G. De, & Gharzouli, R. (2012). Identifying Important Plants Areas (Key Biodiversity Areas for Plants) in northern Algeria. Journal of Threatened Taxa, 4(8), 2753–2765. 10.11609/JoTT.o2998.2753-65

