## Supplementary_materials for "Improving the application of Important Plant Areas to conserve threatened habitats: a case study of Uganda"

### Table S1

Potential natural vegetation types of Uganda (van Breugel et al., 2015) matched alongside the Langdale-Brown et al. (1964) “Vegetation of Uganda”. Matching was supported by unpublished “Vegetation and Climate change in Eastern Africa (VECEA): The case of Uganda”, a precursor to van Breugel et al. (2015). Potential natural vegetation types that were only very marginally within Uganda (Somalia-Masai semi-desert grassland and shrubland – 0.26 km^2^, Acacia-Commiphora stunted bushland – 3.74 km^2^) were removed from this analysis. A plus sign indicates a habitat mosaic between two or more distinct habitat types within either map. Mosaic PNV habitat types were separated out into their constituent habitats for assessment by dividing the area at each time step equally between each habitat type.

| Potential natural vegetation | Langdale-Brown et al. (1964) habitats |
| --- | --- |
| Afromontane undifferentiated forest | Juniperus-Podocarpus Dry Montane Forest; Juniperus-Podocarpus Dry Montane Forest+Acacia-Combretum Savanna; Juniperus-Podocarpus Dry Montane Forest+Combretum-Acacia-Themeda Savanna |
| Dry combretum wooded grassland | Acacia-Albizia-Cenchrus (Beckeropsis)-Cymbopogon+Combretum-Hyparrhenia; Acacia-Albizia-Panicum-Chloris+Combretum-Acacia-Hyparrhenia Savanna; Acacia-Combretum Savanna; Acacia-Combretum Savanna+Acacia-Albizia-Dichrostachys Bushland; Acacia-Ozoroa (Heeria)-Termialia+Commiphora-Euphorbia-Lannea Bushland+Undifferentiated Deciduous Thicket; Acacia-Ozoroa (Heeria)-Terminalia Savanna; Acacia-Ozoroa (Heeria)-Terminalia Savanna+Combretum-Acacia-Lasiurus Savanna; Acacia-Ozoroa (Heeria)-Terminalia+Acacia-Albizia-Dichrostachys Bushland; Albizia-Markhamia Forest+Combretum-Cymbopogon Savanna; Albizia-Markhamia Forest+Combretum-Terminalia-Loudetia; Boswellia-**Zanthoxylum** (Fagara)-Ozoroa (Heeria); Combretum-Acacia-Commiphora; Combretum-Acacia-Commiphora Savanna+Acacia-Commiphora-Lannea Bushland; Combretum-Acacia-Heteropogon; Combretum-Acacia-Heteropogon Savanna+Undifferentiated Deciduous Thicket; Combretum-Acacia-Hyparrhenia; Combretum-Acacia-Hyparrhenia Savanna+Acacia-Themeda-Setaria Savanna; Combretum-Acacia-Hyparrhenia Savanna+Dry Hyparrhenia Grass Savanna; Combretum-Acacia-Hyparrhenia+Acacia-Albizia-Dichrostachys Bushland; Combretum-Acacia-Hyparrhenia+Acacia-Ozoroa (Heeria)-Terminalia+Acacia-Albizia-Dichrostachys Bushland; Combretum-Acacia-Lasiurus; Combretum-Acacia+Acacia-Combretum Savanna; Combretum-Cymbopogon; Combretum-Cymbopogon+Acacia-Themeda-Setaria Savanna; Combretum-Cymbopogon+Moist Hyparrhenia Grass Savanna; Combretum-Hyparrhenia; Combretum-Hyparrhenia Savanna+Combretum-Acacia-Hyparrhenia Savanna; Combretum-Hyparrhenia+Undifferentiated Deciduous Thicket; Combretum-Terminalia-Loudetia; Combretum-Terminalia-Loudetia+Combretum-Acacia-Hyparrhenia Savanna; Combretum-Terminalia-Loudetia+Combretum-Cymbopogon Savanna; Combretum-Terminalia-Loudetia+Combretum-Hyparrhenia; Combretum-Terminalia-Loudetia+Eragrostis-Chloris-Hyparrhenia; Forest/Savanna Mosaic at Medium Altitude+Combretum-Cymbopogon; Forest/Savanna Mosaic at Medium Altitude+Combretum-Terminalia-Loudetia; Juniperus-Podocarpus Dry Montane Forest+Acacia-Combretum Savanna; Lannea-Combretum-Philenoptera (Lonchocarpus); Lannea-Combretum-Philenoptera (Lonchocarpus) Savanna+Acacia-Albizia-Dichrostachys Bushland; Lannea-Combretum-Philenoptera (Lonchocarpus) Savanna+Acacia-Themeda-Setaria Savanna; Lannea-Combretum-Philenoptera (Lonchocarpus) Savanna+Dry Hyparrhenian Grass Savanna; Moist Combretum Savanna+Combretum-Acacia-Hyparrhenia; Moist Combretum Savanna+Combretum-Hyparrrhenia; Undifferentiated Semi-Deciduous Thicket+Albizia Markhamia Forest+Combretum-Cymbopogon Savanna |
| Somalia-Masai Acacia-Commiphora deciduous bushland and thicket | Acacia-Commiphora Bushland; Acacia-Commiphora Bushland+Acacia reficiens-Commiphora Bushland and Thicket; Acacia-Commiphora Thicket; Acacia-Commiphora Thicket+Acacia mellifera Thicket; Acacia-Euphorbia Thicket; Acacia-Lannea Bushland+Acacia-Commiphora Thicket; Acacia-Lannea Bushland+Acacia-Commiphora Thicket+Acacia nubica Thicket; Acacia-Lannea Bushland+Acacia mellifera Thicket; Acacia-Lannea Bushland+Acacia nubica Thicket; Acacia-Lannea Bushland+Commiphora-Euphorbia-Lannea Bushland; Acacia-Lannea Bushland+Undifferentiated Deciduous Thicket; Acacia-mellifera Thicket/Acacia-Setaria Savanna; Acacia mellifera Thicket; Acacia reficiens-Commiphora Bushland and Thicket; Acacia Reficiens-Commiphora Bushland and Thicket+Acacia mellifera Thicket; Acacia seyal-Acacia nilotica-Pennisetum mezianum Bushland+Acacia-Commiphora Thicket; Acacia seyal-Acacia nilotica-Pennisetum mezianum Bushland+Acacia mellifera Thicket; Acacia Tree and Shrub Steppe; Acacia Tree and Shrub Steppe +Acacia-Lannea Bushland; Acacia Tree and Shrub Steppe+Acacia-Albizia-Dichrostachys Bushland; Acacia Tree and Shrub Steppe+Acacia-Lannea Bushland+Acacia mellifera Thicket; Acacia Tree and Shrub Steppe+Acacia-Setaria Savanna; Acacia Tree and Shrub Steppe+Acacia mellifera Thicket; Acacia Tree and Shrub Steppe+Acacia seyal-Acacia nilotica-Pennisetum mezianum Bushland; Commiphora-Euphorbia-Lannea Bushland; Commiphora-Euphorbia-Lannea Bushland+Undifferentiated Deciduous Thicket; Lannea-Acacia-Balanites Bushland+Acacia-Setaria Savanna; Lannea-Acacia-Balanites Bushland+Riparian Thicket+Undifferentiated Deciduous Thicket; Lannea-Acacia-Balanites Bushland+Undifferentiated Deciduous Thicket; Lannea-Acacia Tree and Shrub Steppe; Lannea-Acacia Tree and Shrub Steppe+Lannea-Acacia-Balanites Bushland; Lannea-Acacia Tree and Shrub SteppeAcacia-Commiphora Thicket; Undifferentiated Deciduous Thicket |
| Transition zone of dry Combretum wooded grassland and edaphic wooded grassland on drainage-impeded or seasonally flooded soils | Acacia-Combretum+Acacia-Setaria Savanna; Lannea-Combretum-Philenoptera (Lonchocarpus) Savanna+Acacia-Setaria Savanna |
| Palm wooded grassland | Borassus-Hyperthelia dissoluta (Hyparrhenia dissoluta); Borassus-Hyparrhenia rufa; Borassus-Hyparrhenia rufa+Borassus-Hyperthelia dissoluta (Hyparrhenia dissoluta); Borassus-Hyparrhenia rufa+Undifferentiated Deciduous Thicket; Moist Combretum Savanna+Borassus-Hyperthelia dissoluta (Hyparrhenia dissoluta); Moist Combretum Savanna+Borassus-Hyparrhenia rufa |
| Edaphic grassland and wooded grassland on drainage-impeded or seasonally flooded soils | Acacia-Albizia-Dichrostachys Bushland+Acacia-Setaria Savanna; Acacia-Combretum+Acacia-Setaria Savanna; Acacia-Ozoroa (Heeria)-Terminalia Savanna+Acacia mellifera Bushland+Acacia-Setaria Sananna; Acacia-Imperata Savanna; Acacia-mellifera Thicket/Acacia-Setaria Savanna; Acacia-Setaria Savanna; Acacia-Themeda-Setaria Savanna+Acacia-Setaria Savanna; Acacia-Themeda Savanna; Acacia Tree and Shrub Steppe+Acacia-Setaria Savanna; Borassus-Hyperthelia dissoluta (Hyparrhenia dissoluta)+Themeda-Heteropogon Grass Savanna+Acacia-Imperata Savanna; Borassus-Hyperthelia dissoluta (Hyparrhenia dissoluta)+Sorghastrum Grassland; Brachiaria-Hyparrhenia Grassland; Combretum-Acacia-Hyparrhenia Savanna; Cynometra-Celtis Forest+Acacia-Imperata Savanna; Dry Hyparrhenia Grass Savanna+Acacia Setaria Savanna; Echinochloa Grassland; Echinochloa Grassland+Acacia-Imperata Savanna; Echinochloa Grassland+Acacia-Setaria Savanna; Echinochloa Grassland+Acacia-Themeda Savanna; Echinochloa Grassland+Combretum-Acacia-Hyparrhenia Savanna; Echinochloa Grassland+Cyperus papyrus Swamp; Echinochloa Grassland+Miscanthus (Miscanthidium) Swamp; Echinochloa Grassland+Sorghastrum Grassland; Eragrostis-Loudetia Grass Savanna+Sorghastrum Grassland; Eragrostis-Loudetia Grass Savanna; Forest/Savanna Mosaic at Medium Altitude+Eragrostis-Loudetia Grass Savanna; Lannea-Acacia-Balanites Bushland+Acacia-Setaria Savanna; Lannea-Acacia-Balanites Bushland+Riparian Thicket+Undifferentiated Deciduous Thicket; Lannea-Combretum-Philenoptera (Lonchocarpus) Savanna+Acacia-Setaria Savanna; Piptadeniastrum-Albizia-Celtis Forest+Acacia-Imperata Savanna; Riparian Thicket; Riparian Thicket+Acacia-Setaria Savanna; Riparian Thicket+Acacia-Themeda-Setaria Savanna+Acacia-Setaria Savanna; Sorghastrum Grassland; Sorghastrum Grassland+Acacia-Imperata Savanna; Sorghastrum Grassland+Combretum-Acacia-Hyparrhenia Savanna; Themeda-Heteropogon Grass Savanna+Acacia-Imperata Savanna; Undifferentiated Deciduous Thicket+Combretum-Acacia-Hyparrhenia Savanna; Undifferentiated Semi-Deciduous Thicket+Acacia-Themeda-Setaria Savanna |
| Vitellaria (Butyrospermum) wooded grassland | Vitellaria (Butyrospermum)-Daniellia-Hyparrhenia; Vitellaria (Butyrospermum)-Hyperthelia dissoluta (Hyparrhenia dissoluta); Vitellaria (Butyrospermum)-Hyparrhenia rufa; Vitellaria (Butyrospermum)-Hyparrhenia rufa+Vitellaria (Butyrospermum)-Hyperthelia dissoluta (Hyparrhenia dissoluta); Moist Combretum Savanna+ Vitellaria (Butyrospermum)-Hyparrhenia rufa; Undifferentiated Semi-Deciduous Thicket+Vitellaria (Butyrospermum)-Hyperthelia dissoluta (Hyparrhenia dissoluta) |
| Evergreen and semi-evergreen bushland and thicket | Albizia-Markhamia Forest+Undifferentiated Semi-Deciduous Thicket; Celtis-Gambeya (Chrysophyllum) Forest+Undifferentiated Semi-Deciduous Thicket; Celtis-Gambeya (Chrysophyllum) Forest+Undifferentiated Semi-Deciduous Thicket+Moist Hyparrhenia Grass Savanna; Undifferented Semi-Deciduous Thicket; Undifferentiated Semi-Deciduous Thicket+Albizia Markhamia Forest+Combretum-Cymbopogon Savanna; Undifferentiated Semi-Deciduous Thicket+Themeda Heteropogon Grass Savanna; Undifferentiated Semi-Deciduous Thicket+Themeda Loudetia Grass Savanna |
| Upland Acacia wooded grassland | Acacia-Themeda-Setaria Savanna |
| Lowland bamboo | Lowland Bamboo Thicket |
| Transitional zone Somalia-Masai Acacia-Commiphora deciduous bushland and thicket and Dry Combretum wooded grassland | Acacia-Combretum Savanna+Acacia-Albizia-Dichrostachys Bushland; Acacia-Ozoroa (Heeria)-Termialia+Commiphora-Euphorbia-Lannea Bushland+Undifferentiated Deciduous Thicket; Acacia-Ozoroa (Heeria)-Terminalia Savanna+Acacia mellifera Bushland+Acacia-Setaria Sananna; Acacia-Ozoroa (Heeria)-Terminalia+Acacia-Albizia-Dichrostachys Bushland; Combretum-Acacia-Commiphora Savanna+Acacia-Commiphora-Lannea Bushland; Combretum-Acacia-Hyparrhenia+Acacia-Albizia-Dichrostachys Bushland; Combretum-Acacia-Hyparrhenia+Acacia-Ozoroa (Heeria)-Terminalia+Acacia-Albizia-Dichrostachys Bushland; Combretum-Acacia-Lasiurus+Combretum-Acacia-Commiphora+Acacia-Commiphora Bushland; Lannea-Combretum-Philenoptera (Lonchocarpus) Savanna+Acacia-Albizia-Dichrostachys Bushland; Moist Combretum Savanna+Combretum-Hyparrrhenia |
| Freshwater swamp | Cyperus papyrus Swamp; Cyperus papyrus Swamp+Miscanthus (Miscanthidium) Swamp; Miscanthus (Miscanthidium) Swamp |
| Riverine wooded vegetation | Riparian Thicket; Undifferentiated Semi-Deciduous Thicket+Acacia-Themeda-Setaria Savanna |
| Moist Combretum wooded grassland | Albizia-Combretum Woodland; Moist Combretum Savanna |
| Lake Victoria drier peripheral semi-evergreen Guineo-Congolian rain forest | Albiza-Milicia (Chlorophora) Forest; Albizia-Milicia (Chlorophora) Forest+Undifferentiated Semi-Deciduous Thicket; Albizia-Milicia (Chlorophora) Forest+Undifferentiated Semi-Deciduous Thicket+Moist Combretum Savanna; Celtis-Gambeya (Chrysophyllum) Forest; Celtis-Gambeya (Chrysophyllum) Forest+Undifferentiated Semi-Deciduous Thicket; Celtis-Gambeya (Chrysophyllum) Forest+Undifferentiated Semi-Deciduous Thicket+Moist Hyparrhenia Grass Savanna; Cynometra-Celtis Forest; Cynometra-Celtis Forest+Acacia-Imperata Savanna; Cynometra-Celtis Forest+Moist Combretum Savanna; Parinari Forest; Piptadeniastrum-Albizia-Celtis Forest; Piptadeniastrum-Albizia-Celtis Forest+Acacia-Imperata Savanna; Piptadeniastrum-Uapaca Forest; Piptadeniastrum-Uapaca Forest+Piptadeniastrum-Albizia-Celtis Forest; Piptadeniastrum-Uapaca Forest+Themeda-Loudetia Grass Savanna |
| Climatic grasslands | Grass Steppe; Grass Steppe+Acacia seyal-Acacia nilotica-Pennisetum mezianum Bushland |
| Vitex-Phyllanthus-Shirakiopsis (Sapium)-Terminalia and Terminalia glaucescens woodland | Terminia Woodland; Vitex-Phyllanthus-Shirakiopsis (Sapium)-Terminalia Woodland; Vitex-Phyllanthus-Shirakiopsis (Sapium)-Terminalia Woodland+Moist Combretum Savanna; Vitex-Phyllanthus-Shirakiopsis (Sapium)-Terminalia Woodland+Terminalia Woodland |
| Edaphic wooded grassland on drainage-impeded or seasonally flooded soils or riverine wooded vegetation | Riparian Thicket+Acacia-Setaria Savanna; Riparian Thicket+Acacia-Themeda-Setaria Savanna+Acacia-Setaria Savanna |
| Edaphic grassland on drainage-impeded or seasonally flooded soils and palm wooded grassland | Borassus-Hyperthelia dissoluta (Hyparrhenia dissoluta)+Sorghastrum Grassland |
| Dry Combretum wooded grassland + palm wooded grassland | Borassus-Hyperthelia dissoluta (Hyparrhenia dissoluta)+Combretum-Hyparrhenia |
| Evergreen and semi-evergreen bushland + thicket and palm wooded grassland | Borassus-Hyparrhenia rufa+Undifferentiated Deciduous Thicket |
| Montane Ericaceous belt | Ericaceae-Seriphium (Stoebe) Heath |
| Afromontane rain forest | Forest/Savanna Mosaic at High Altitude+Montane Thicket; Montane Thicket; Prunus (Pygeum) Moist Montane Forest |
| Afromontane Bamboo and Hagenia | Hagenia-Rapanea Moist Montane Forest; Hagenia-Rapanea Moist Montane Forest+Oldeania (Arundinaria) Montane Bamboo Forest |
| Edaphic wooded grassland on drainage-impeded or seasonally flooded soils + palm wooded grassland | Borassus-Hyperthelia dissoluta (Hyparrhenia dissoluta)+Themeda-Heteropogon Grass Savanna+Acacia-Imperata Savanna |
| Afroalpine vegetation | Alchemilla-Helichrysum Moorland |
| Swamp forest | Baikiaea-Podocarpus Seasonal Swamp Forest; Rauvolfia-Croton Seasonal Swamp Forest |
| Vitellaria-Combretum Mosaic | Vitellaria (Butyrospermum)-Hyperthelia dissoluta (Hyparrhenia dissoluta)+Combretum-Acacia-Hyparrhenia; Vitellaria (Butyrospermum)-Hyperthelia dissoluta (Hyparrhenia dissoluta)+Combretum-Hyparrhenia |

### Table S2

NFA Land Use Cover (NFA 2017) map classified into natural and non-natural vegetation types

| LUC code | LUC class land cover types | Natural |
| --- | --- | --- |
| 1 | Broadleaved plantation | No |
| 2 | Coniferous Plantation | No |
| 3 | THF well stocked | Yes |
| 4 | THF low stocked | Yes |
| 5 | Woodland | Yes |
| 6 | Bushland | Yes |
| 7 | Grassland | Yes |
| 8 | Wetland | Yes |
| 9 | Subsistence farmland | No |
| 10 | Commercial farmland | No |
| 11 | Built up area | No |
| 12 | Water bodies | Yes |
| 13 | Impediments | Yes |
